# Refinement of protein structure via contact based potentials in replica exchange simulations

**DOI:** 10.1101/2020.02.14.949172

**Authors:** Arthur Voronin, Marie Weiel, Alexander Schug

## Abstract

Proteins are complex biomolecules which perform critical tasks in living organisms. Knowledge of a protein’s structure is essential for understanding its physiological function in detail. Despite the incredible progress in experimental techniques, protein structure determination is still expensive, time-consuming, and arduous. That is why computer simulations are often used to complement or interpret experimental data. Here, we explore how *in silico* protein structure determination based on replica exchange molecular dynamics can benefit from including contact information derived from theoretical and experimental sources, such as direct coupling analysis or NMR spectroscopy. To reflect the influence from erroneous and noisy data we probe how false-positive contacts influence the simulated ensemble. Specifically, we integrate varying numbers of randomly selected native and non-native contacts and explore how such a bias can guide simulations towards the native state. We investigate the number of contacts needed for a significant enrichment of native-like conformations and show the capabilities and limitations of this method. Adhering to a threshold of approximately 75% true-positive contacts within a simulation, we obtain an ensemble with native-like conformations of high quality. We find that contact-guided REX MD is capable of delivering physically reasonable models of a protein’s structure.

**Author summary:** Protein structure prediction, that is obtaining a protein structure starting from a sequence using any computational method, is a great challenge. Over the past years a broad variety of methods evolved, ranging from algorithms for “blind” or *de novo* predictions using Monte-Carlo or physics-based biomolecular simulation methods to algorithms transferring structure information obtained from known homologous proteins. Recently, purely data-driven approaches using neural networks have shown to be capable of predicting high-quality structures. However, some local structural motifs are only poorly resolved and need further refinement. Here, we explore to what extent contact information helps guiding replica exchange molecular dynamics towards the native fold. By adding a contact pair bias potential to the energy function, we effectively guide the search towards the target structure by narrowing the conformational space to be sampled. We find that such an energetic bias, even if containing false-positive contacts to a certain extent, greatly enhances the refinement process and improves the chance of finding the native state in a single run.

## Introduction

Knowledge of protein structures is crucial for understanding the proteins’ functions or the biological processes they take part in. Structural knowledge is also critical in related fields such as pharmacology to design drugs or medicine to understand the molecular origins of pathogenesis. Both protein structure and function are intrinsically encoded in the corresponding amino acid sequence [1–3]. Over the past years, experimental techniques to gain such sequential data have become exceptionally efficient and lead to fast growing sequence databases, e.g., GenBank [4] and UniProt [5]. In contrast, experimental structure determination with methods such as X-ray crystallography or NMR spectroscopy are comparably time-consuming and expensive with often involved procedures. Some other experimental techniques, e.g., SAXS, FRET, or cryoEM, do not directly provide structural data but have to be carefully interpreted [6–8].

Computer simulations are commonly used to interpret and complement experimental data. Novel, purely data-driven approaches can predict protein structures of high quality [9, 10] but lack insight into the physical processes driving structure adoption and cannot be easily complemented by experimental information. Depending on the method, local structural motifs are often less resolved [9] and could benefit from additional refinement. Physics-driven approaches are based on energy functions called force fields. Lindorff et al. demonstrated by Molecular Dynamics (MD) simulations that current force fields are sufficiently accurate to reversibly fold proteins starting from unfolded conformations [11, 12]. Still, the computational cost for such *de novo* folding simulations is extremely high. As a result, simulations on the millisecond timescale can currently only be performed on specialized supercomputers like Anton. Luckily, it is possible to guide the simulations towards target structures or ensembles by introducing an energetic bias based on experimental data. This bias helps smoothing the energy landscape, which has frustrated, glassy properties with many competing minima separated by high barriers. At the same time, the computational costs are lowered due to the reduced sampling space. One can further lower computational demands by enhanced sampling techniques [13–15].

In this work, we assume having access to varying amounts of error-ridden contact information as an additional potential bias for MD simulations. Such contact information, i.e. information about adjacent amino acids, can be obtained from different sources. Sparse NMR contact maps would be one example. By themselves they provide insufficient information for structure generation and have to be complemented. Recently, contacts derived from coevolution analysis methods such as direct coupling analysis (DCA) [16] infer contact information from large multiple sequence alignments. DCA identifies coevolving residue pairs, which can be interpreted as spatially adjacent. This information was successfully used for structure prediction [17] even in large-scale studies of proteins [18] or for RNA [19]. However, it is often uncertain how error-prone contact information is. NMR assignments can be wrong or DCA can contain false-positive contacts. For this purpose, we performed an extensive study to investigate the influence of native (“correct”) and non-native (“wrong”) contact information with regards to structure refinement. Additionally, to overcome residual entrapment and the occurring multiple-minima problem during the simulation process, we use replica exchange (REX) as an enhanced sampling technique [11, 20–22].

By combining both contact information and REX MD simulations, we drastically enrich native or native-like conformations in the simulated ensemble of a single run. To systematically study and test our method’s performance, we investigate two small proteins with known native structures. Starting from an unfolded state, we look at contact-guided structure determination with REX MD simulations. We test different scenarios (see Table 1) by varying the bias quality, i.e. changing the true-positive rate (TPR) and the total number of randomly selected contact pairs. We intentionally apply an equal force coefficient *k* (see Eq 7) to all used contacts, as the study is performed to estimate the influence resulting from both native and non-native contacts. We analyze the data for each test case with simulated times of 250 ns, especially for the lowest temperature replica. By comparing the test cases to a reference simulation not including any contact information, we can thus estimate the total number of required restraints and the bias strength. The study shows good results for both tested proteins as long as the used bias quality was above a certain threshold. It is possible to include additional experimental bias into such simulations and use them as a tool for hybrid data integration.

**Table 1.** Variation of bias quality during method performance study using REX simulations. Overview of the fourteen tested scenarios during the method performance study which were conducted on both test proteins. Listed are the true positive rate (TPR) of used contact pairs (CP) in percent, number of CP used as restraints, number of native contacts and number of non-native contacts. The coloring is used to highlight the respective contacts of the proteins (cf. SI).

| TPR (%) | ref | 100 | 100 | 100 | 100 | 100 | 75 | 75 | 75 | 75 | 50 | 50 | 50 | 50 |
| --- | --- | --- | --- | --- | --- | --- | --- | --- | --- | --- | --- | --- | --- | --- |
| # CP | 0 | 6 | 12 | 24 | 36 | 48 | 12 | 24 | 36 | 48 | 12 | 24 | 36 | 48 |
| # native | 0 | 6 | 12 | 24 | 36 | 48 | 9 | 18 | 27 | 36 | 6 | 12 | 18 | 24 |
| # non-native | 0 | 0 | 0 | 0 | 0 | 0 | 3 | 6 | 9 | 12 | 6 | 12 | 18 | 24 |
| # Color | - | blue | cyan | green | orange | red | cyan | green | orange | red | cyan | green | orange | red |

## Results and discussion

### Test systems

We use two well-known proteins for our systematic study. The first candidate is the 20-residue miniprotein Trp-Cage (PDB: 1l2y [23]). Its tertiary structure consists of an *α*-helix followed by a turn and a 3/10-helix. Trp-Cage was specifically designed as a fast folder and reaches folding timescales of approximately 4 *µ*s [24]. The second test system is Villin Headpiece (VHP, PDB: 1vii [25]). This protein has a sequence length of 35 residues and forms a three-helix structure. Similar to Trp-Cage, VHP can also achieve folding times in the order of *µ*s [26, 27].

### Trp-Cage

We performed REX simulations of Trp-Cage for a total of 60 replica yielding trajectories of 250 ns simulated time ranging from *T*_0_= 300*K* to *T*_59_ = 624.67*K*. To compare the different scenarios as listed in Table 2, each REX simulation initiates from the same unfolded conformation. As given in Eq (7), a sigmoid potential with the same coupling strength *k* is assigned to all implemented restraints (only C_*α*_-C_*α*_ pairs). Note that the resulting force is distance-dependent and the potential has a limit of 10 kJ mol^−1^. As a first analysis step, we checked if the coupling strength of the implemented restraints is adequate. The energetic bias is supposed to guide the protein towards a native-like structure. We determined the time-dependent backbone RMSD with respect to the native structure for all replica in each test case. To get an overview and quickly compare the RMSD statistics across all scenarios, the color-coded RMSD values of all replica are displayed in heatmap plots. Fig 1A and Fig 1B exemplary show a comparison between the reference simulation and the simulation with 12 native contacts, respectively.

**Table 2.** Total Score (TS) and High Accuracy (HA) percentiles of Trp-Cage REX simulations. Overview of percentile values at lowest temperature replica. Listed are the true-positive rate (TPR) in percent, number of used contact pairs (CP) as restraints, Global Distance Test Total Score percentiles denoted as *P*^TS^, and Global Distance Test High Accuracy percentiles denoted as *P*^HA^. Values equal to or greater than the reference values are shaded in gray. According to Eq (2) significantly greater values are bold.

| TPR (%) | # CP | $P_{80}^{TS}$ | $P_{85}^{TS}$ | $P_{90}^{TS}$ | $P_{95}^{TS}$ | $P_{100}^{TS}$ | $P_{80}^{HA}$ | $P_{85}^{HA}$ | $P_{90}^{HA}$ | $P_{95}^{HA}$ | $P_{100}^{HA}$ |
| --- | --- | --- | --- | --- | --- | --- | --- | --- | --- | --- | --- |
| ref | 0 | 53.75 | 88.75 | 93.75 | 96.25 | 100.00 | 30.00 | 67.50 | 76.25 | 81.25 | 98.75 |
| 100 | 6 | **96.25** | **96.25** | **97.50** | 97.50 | **100.00** | **80.00** | 81.25 | 82.50 | 85.00 | 96.25 |
| 100 | 12 | **96.25** | **97.50** | **97.50** | **98.75** | **100.00** | **81.25** | 82.50 | 83.75 | 86.25 | 97.50 |
| 100 | 24 | **97.50** | **97.50** | **97.50** | **98.75** | **100.00** | **82.50** | **83.75** | 85.00 | 86.25 | 97.50 |
| 100 | 36 | **97.50** | **97.50** | **97.50** | **98.75** | **100.00** | **82.50** | **83.75** | 85.00 | 86.25 | 97.50 |
| 100 | 48 | **97.50** | **97.50** | **98.75** | **98.75** | **100.00** | **82.50** | **83.75** | 85.00 | 86.25 | 97.50 |
| 75 | 12 | **95.00** | **96.25** | **97.50** | 97.50 | **100.00** | **78.75** | 80.00 | 82.50 | 85.00 | **98.75** |
| 75 | 24 | **95.00** | **96.25** | 96.25 | 97.50 | **100.00** | **77.50** | 80.00 | 81.25 | 83.75 | 96.25 |
| 75 | 36 | **96.25** | **96.25** | **97.50** | 97.50 | **100.00** | **80.00** | 81.25 | 82.50 | 85.00 | 96.25 |
| 75 | 48 | **96.25** | **96.25** | **97.50** | **98.75** | **100.00** | **80.00** | 81.25 | 83.75 | 85.00 | 97.50 |
| 50 | 12 | 41.25 | 47.50 | 85.00 | 95.00 | **100.00** | 16.25 | 23.75 | 62.50 | 77.50 | 96.25 |
| 50 | 24 | 40.00 | 42.50 | 47.50 | 91.25 | **100.00** | 17.50 | 20.00 | 23.75 | 71.25 | 96.25 |
| 50 | 36 | 36.25 | 38.75 | 41.25 | 43.75 | 95.00 | 15.00 | 17.50 | 20.00 | 22.50 | 82.50 |
| 50 | 48 | 37.50 | 38.75 | 42.50 | 46.25 | 93.75 | 15.00 | 17.50 | 18.75 | 23.75 | 76.25 |

**Fig. 1.**
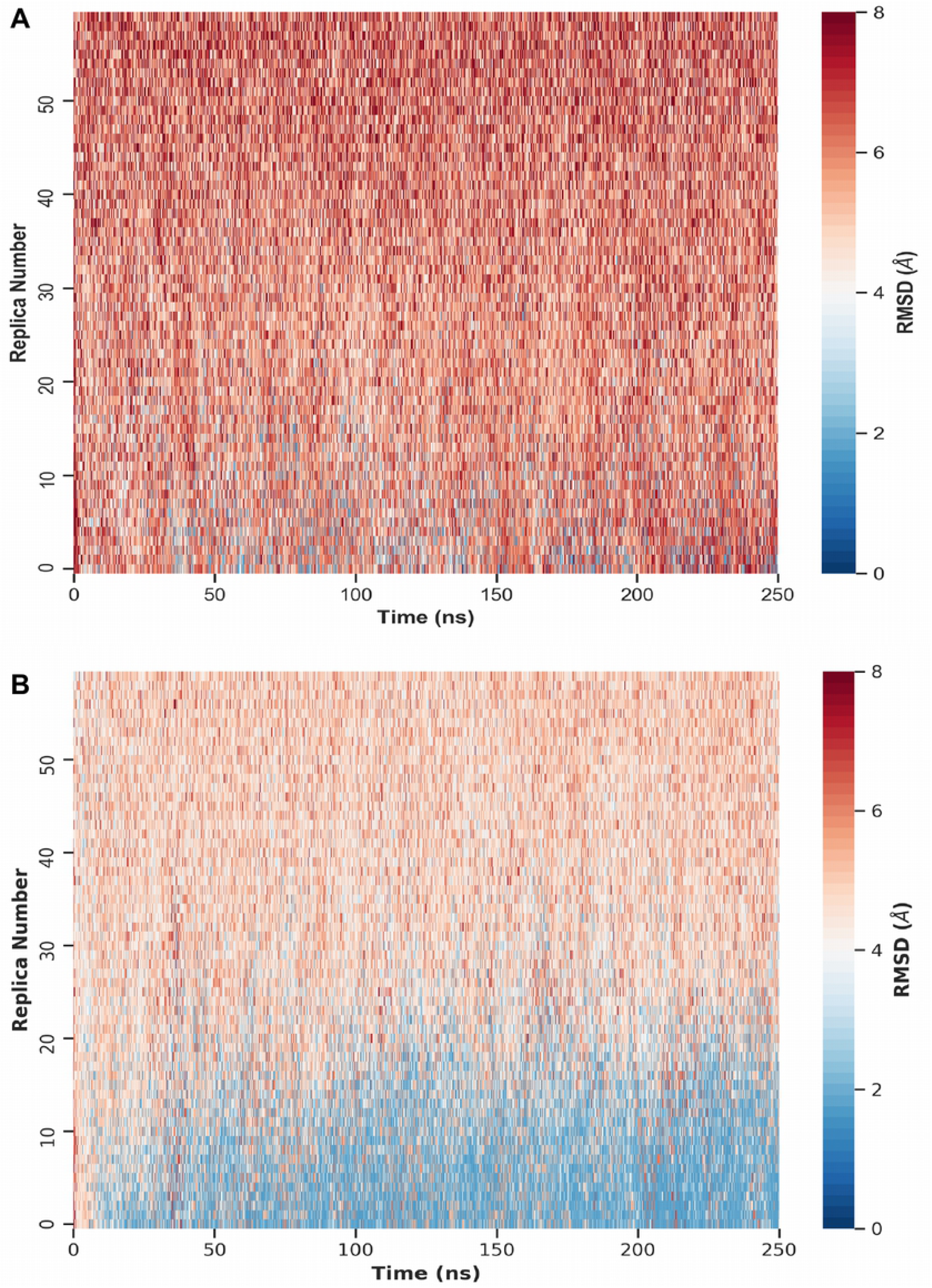
Backbone RMSD time evolution of Trp-Cage REX simulations. (A) Reference simulation without contact information. (B) Simulation with bias of 12 native contacts.

The reference run mostly shows RMSD values greather than 4 Å with a few random exceptions at lower temperature replica throughout the simulation. As expected, introducing a bias potential with purely native contact restraints strongly improves the RMSD values for lower temperature replica, as displayed in Fig 1B. Here, the majority of low-temperature RMSD values turns from red (more than 4 Å) to blue (less than 4 Å) compared to the reference simulation. Heatmap plots of the remaining cases using only ideal contact pairs at 100% TPR can be found in S1 Fig to S3 Fig. They prove that the number of used native contact restraints is correlated with the enrichment of native-like conformations. The blue region therefore grows with increased “correct” bias compared to the unbiased reference simulation. The greatest step-wise improvement for lower temperature replica is observed for the transition from 6 to 12 native contacts. With any sort of contact information usually being error-prone, it is nearly impossible to apply a perfect bias corresponding to a TPR of approximately 100%. Heatmap plots for the other REX simulations with TPRs of 75% and 50% are shown in S4 Fig to S7 Fig. Mixed scenarios containing both true- and false-positive contact information are performed to estimate the bias quality threshold required to improve structure prediction. For this purpose, we analyzed primarily the lowest temperature replica to see which conformations become enriched. For each tested scenario, a so-called Δ*F* histogram displays the frequency difference of observed backbone RMSD values to the reference. Histograms for Trp-Cage are summarized in Fig 2. Simulations with purely native contacts (Fig 2A to Fig 2D) show a strong enrichment of conformations with RMSD values between 1.6 and 3.0 Å as indicated by the green region. Frequencies of conformations with RMSD values above 3.0 Å got reduced accordingly. In case of Trp-Cage, the net gain of native-like folds does not improve when the bias exceeds 12 restraints, which corresponds to approximately *L/*2 contact pairs. Test cases with mixed contacts at 75% TPR (Fig 2E to Fig 2H) show a similar behavior. Additionally, the two scenarios with 12 and 24 mixed contacts also enrich conformations with RMSDs around 5.0 Å. Such conformations can be filtered out in a subsequent step using a suitable method or algorithm for frame selection. Scenarios with a low bias quality of 50% TPR (Fig 2I to Fig 2L) show worse statistics compared to the reference simulations and are therefore inappropriate for the intended purpose. Naively one might guess that the “correct” bias would cancel the “wrong” bias out. However, if implemented restraints are clustered within the contact map, the local bias adds up and apparently is strong enough to trap the protein in unfavored conformations. When using large numbers of contact restraints with respect to the protein length, it is necessary to reduce the overall force coefficient *k* of the sigmoid potential (see Eq 7) to compensate this effect.

**Fig. 2.**
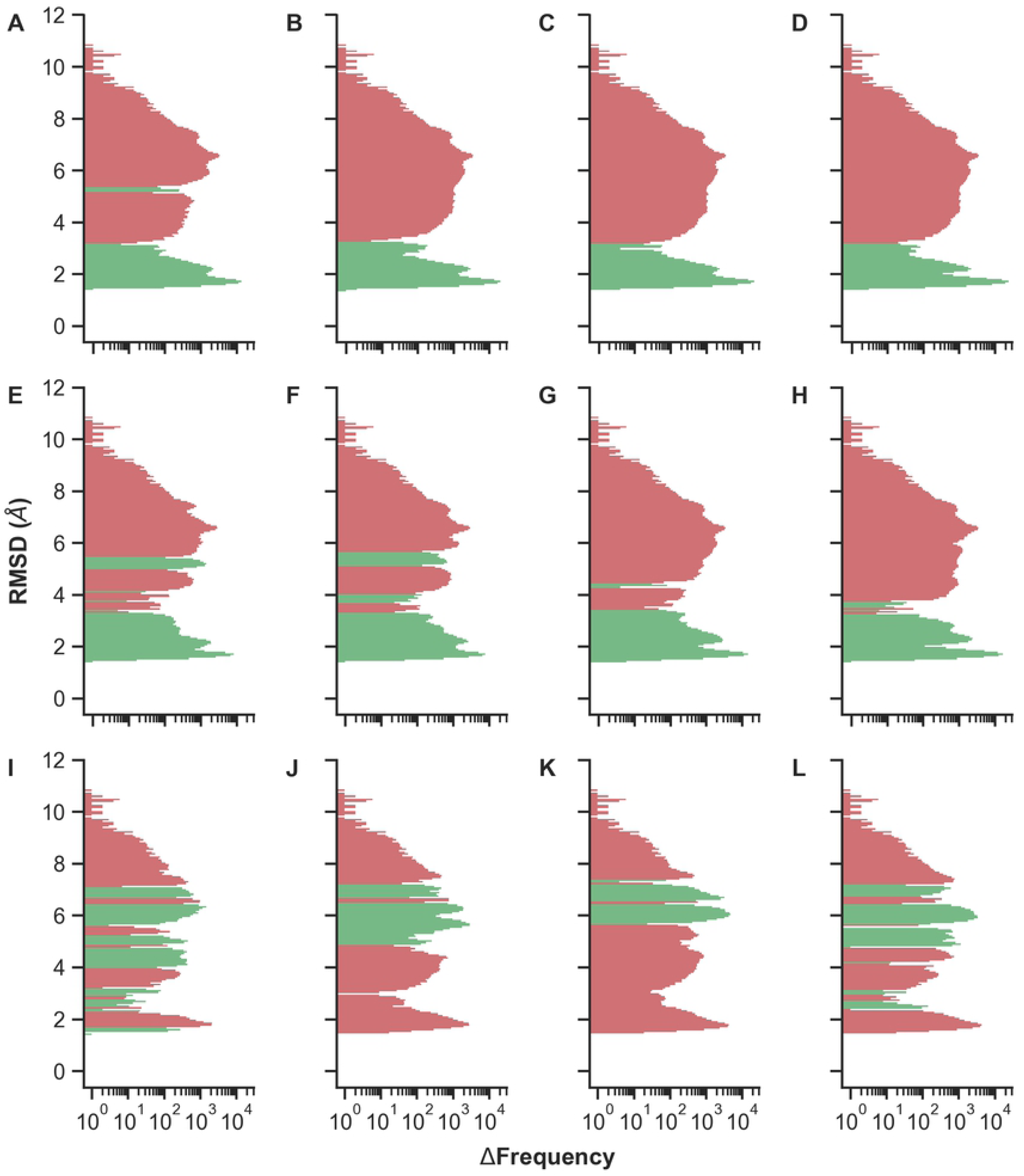
Δ*F* histograms of Trp-Cage REX simulations. Histograms show the enrichment and depletion of conformations with a particular backbone RMSD as compared to the reference. Histogram bins are defined by the RMSD axis, while the logarithmic Δ*F* axis displays Δ*F* = *N*_sim_ − *N*_ref_. Positive (negative) values corresponding to enrichment (depletion) are shown in green (red). (A-D) Simulations with 100% TPR and 12, 24, 36, 48 restraints. (E-H) Simulations with 75% TPR and 12, 24, 36, 48 restraints. (I-L) Simulations with 50% TPR and 12, 24, 36, 48 restraints.

The disadvantage of RMSD-based evaluation of structure quality is that local deviations between mobile and target structure already result in a disproportionate increase. This is why we transition to the so-called Global Distance Test (GDT) which takes local misalignments better into account. The distributions of the two scores *GDT*_TS_ and *GDT*_HA_ provide an additional perspective to the estimation of the bias quality necessary for an effective integration of contact information into REX simulations. Table 2 gives an overview of occurring percentiles of *GDT*_TS_ and *GDT*_HA_ scores for all REX MD simulations with Trp-Cage as a test system. To simplify the comparison, shaded table cells indicate improved percentiles *P*_x_ compared to the reference *P*_x,ref_ for each score variant, i.e. cells with

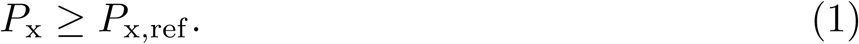

Additionally, we use a bold font for values which satisfy

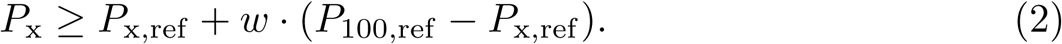

to highlight a significant improvement. In Eq (2) each *P*_x_ is compared to a percentile-specific threshold only depending on corresponding reference values. The threshold is defined as the sum of the percentile itself and a weighted difference of this percentile to the highest observed value. The difference indicates the practically possible improvement in relation to the reference. To determine a “significant” improvement, we set the coefficient *w* to 50%. In scenarios with TPRs of 75% or higher, the TS distribution is shifted significantly from 53.75 to score values above 96 already at the 80th percentile. This means that 20% of the trajectory already consist of conformations almost identical to the known native structure. The table section with HA scores similarly shows a significant improvement. Note that the reference simulation yielded an exceptional HA score of 98.75 during one turnaround of the REX simulation.

The comparison of 50% TPR simulations to the reference shows that highly error-prone contact bias has a very negative influence and is not sufficient to effectively improve structure determination in REX.

### Villin Head Piece

All VHP REX simulations were performed under the same conditions as for Trp-Cage. However, due to the increased system size 40 additional replica were required to achieve nearly constant exchange rates across the considered temperature range. The REX RMSD heatmap plots (see S8 Fig to S14 Fig) have similar but less prominent tendencies compared to Trp-Cage. The reference shows very poor RMSD statistics throughout. Even with enhanced sampling as in REX we observe that the simulated time span of 250 ns is too short for VHP to be guided towards the native structure without additional bias. As soon as the bias potential is activated, we clearly see an analogous growth of the blue region within the heatmap plots indicating enrichment of native-like conformations. Furthermore, the most potent improvement was observed for the transition from 12 to 24 native contacts as shown in S9 Fig. The Δ*F* histograms of the lowest temperature replica are summarized in Fig 3. Fig 3A to Fig 3D illustrate scenarios under perfect conditions, i.e. at a TPR of 100%. In the case of VHP, large improvements are made up to 24 restraints. Scenarios with 36 and 48 restraints show almost identical results as the simulation with 24 restraints. We observe conformation frequencies with RMSDs above 4.0 Å to be reduced while conformations with RMSDs between 2.0 and 4.0 Å got enriched. Scenarios with mixed contacts at 75% TPR are displayed in Fig 3E to Fig 3H. We still observe an enrichment of low RMSD conformations but also a drastic increase of conformations with values around 5.0 to 8.0 Å. This is the result of the first three non-native contacts included, which were randomly selected for this test case. Long-range contact pairs (*i, j*), which are far away from the main diagonal of a contact map, have a more significant influence compared to contact pairs with a small difference in their sequence numbers Δ_*ij*_ = |*i* − *j*|close to the main diagonal. All simulations with a TPR of 75% nonetheless show a net gain of native-like conformations. Therefore, they should be favored over the unbiased scenario given a functional algorithm to filter the best structures. Scenarios with a TPR of 50% are displayed in Fig 3I to Fig 3L. Here, we observe that both low and high RMSD conformations appear less compared to the reference, whereas conformations between 5.0 and 9.0 Å are enriched. The high ratio of non-native contacts lowers the chance of successful structure determination as before for Trp-Cage.

**Fig. 3.**
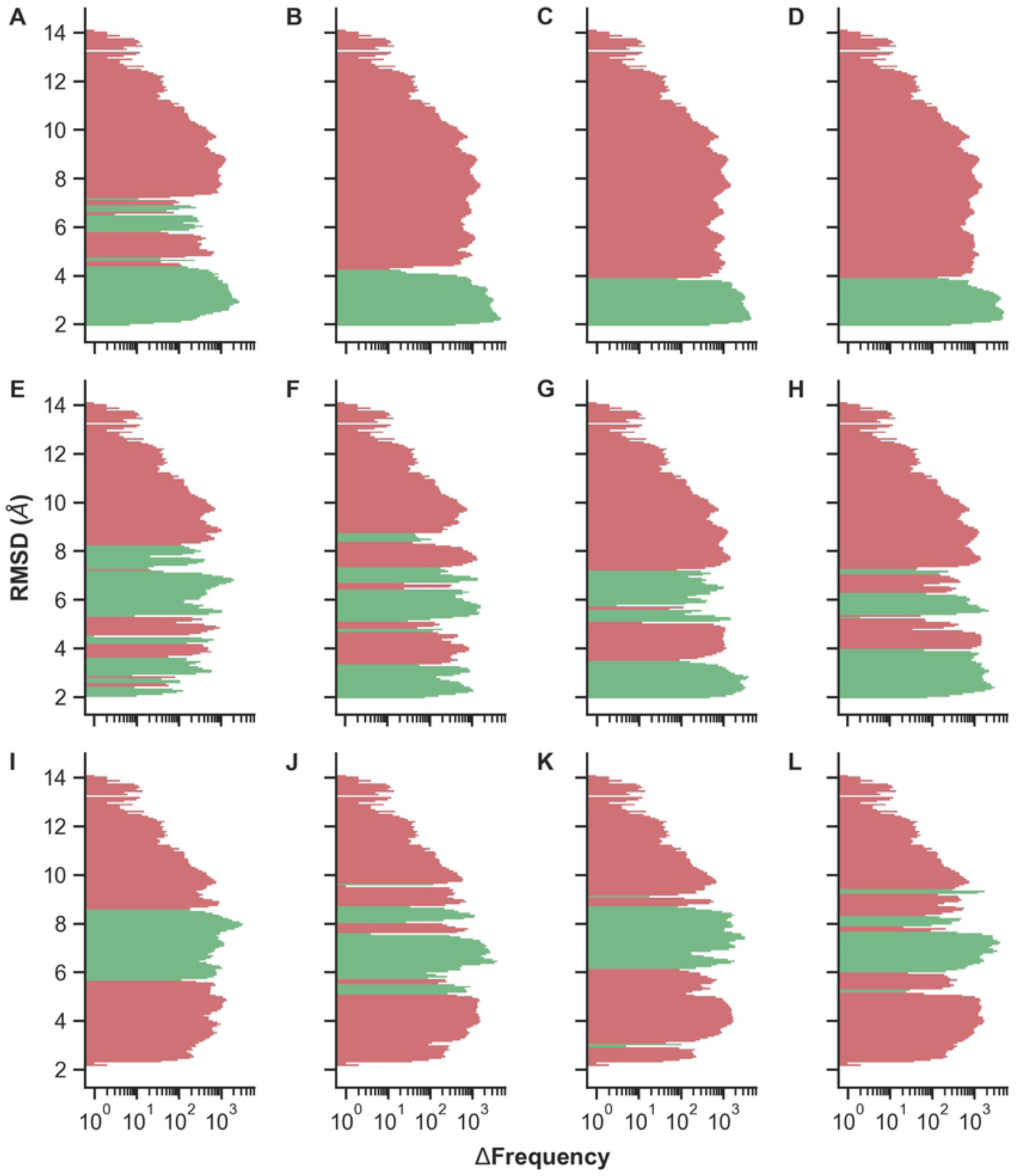
Δ*F* histograms of VHP REX simulations. Histograms show the enrichment and depletion of conformations with a particular backbone RMSD as compared to the reference. Histogram bins are defined by the RMSD axis, while the logarithmic Δ*F* axis displays Δ*F* = *N*_sim_ − *N*_ref_. Positive (negative) values corresponding to enrichment (depletion) are shown in green (red). (A-D) Simulations with 100% TPR and 12, 24, 36, 48 restraints. (E-H) Simulations with 75% TPR and 12, 24, 36, 48 restraints. (I-L) Simulations with 50% TPR and 12, 24, 36, 48 restraints.

Table 3 specifies the percentiles of *GDT* _TS_ and *GDT* _HA_ score distributions. We find that the VHP REX simulations benefitted from restraints with a TRP of 75% or higher, resulting in a significant increase of GDT score statistics. For ideal results in mixed scenarios with 75% TPR, a restraint number in the order of the protein length is required. Analogous to our previous observation during the RMSD-based discussion, simulations with only 50% TPR were worse compared to the reference scenario. We conclude that a bias of such poor quality is therefore unsuited to achieve useful results within contact-guided REX simulations. To highlight the local accuracy and visualize how well the simulated structures fit the native state, Fig 4A gives an overview of the best observed structures during the 75% TPR simulation ranked by HA scores. Each protein residue is assigned a colored rectangle representing the C_*α*_-C_*α*_ distance between mobile and reference after a least-squares fit. As evident in Fig 4A, many simulated structures greatly resemble the target structure which manifests in C_*α*_ displacements below 2 Å. The best observed structure with an HA score of 70.83 corresponding to the first line is displayed in Fig 4B. Local accuracy figures with 36 restraints and varying TPR can be looked up in S16 Fig to S18 Fig.

**Table 3.** Total Score (TS) and High Accuracy (HA) percentiles of VHP REX simulations. Overview of percentile values at lowest temperature replica. Listed are the true-positive rate (TPR) in percent, number of used contact pairs (CP) as restraints, Global Distance Test Total Score percentiles denoted as *P*^TS^, and Global Distance Test High Accuracy percentiles denoted as *P*^HA^. Values equal to or greater than the reference values are shaded in gray. According to Eq (2) significantly greater values are bold.

| TPR (%) | # CP | $P_{80}^{TS}$ | $P_{85}^{TS}$ | $P_{90}^{TS}$ | $P_{95}^{TS}$ | $P_{100}^{TS}$ | $P_{80}^{HA}$ | $P_{85}^{HA}$ | $P_{90}^{HA}$ | $P_{95}^{HA}$ | $P_{100}^{HA}$ |
| --- | --- | --- | --- | --- | --- | --- | --- | --- | --- | --- | --- |
| ref | 0 | 50.00 | 53.47 | 57.64 | 63.89 | 79.17 | 27.08 | 30.56 | 34.72 | 40.98 | 58.34 |
| 100 | 6 | **66.67** | **68.75** | **71.53** | **75.00** | **87.50** | **43.06** | **45.14** | **47.92** | **51.39** | **68.06** |
| 100 | 12 | 61.11 | 63.19 | 65.97 | 69.44 | **86.11** | 37.50 | 39.58 | 42.36 | 45.84 | **65.28** |
| 100 | 24 | **71.53** | **73.61** | **75.00** | **77.08** | **88.89** | **47.92** | **49.30** | **51.39** | **53.47** | **68.75** |
| 100 | 36 | **71.53** | **73.61** | **75.00** | **77.08** | **88.89** | **47.92** | **50.00** | **51.39** | **54.16** | **70.83** |
| 100 | 48 | **72.22** | **73.61** | **75.00** | **77.08** | **87.50** | **48.61** | **50.00** | **51.39** | **53.47** | **68.06** |
| 75 | 12 | 47.92 | 50.70 | 54.17 | 59.02 | **87.50** | 24.30 | 27.08 | 30.56 | 34.72 | **65.97** |
| 75 | 24 | 49.30 | 54.86 | 59.02 | 70.14 | **84.72** | 25.00 | 30.56 | 34.72 | 45.83 | **64.58** |
| 75 | 36 | **68.06** | **71.53** | **74.31** | **77.08** | **88.89** | **43.75** | **47.22** | **50.00** | **53.47** | **70.83** |
| 75 | 48 | 62.50 | 65.97 | **69.44** | **73.61** | **85.42** | 38.89 | 42.36 | 45.83 | 49.30 | **63.89** |
| 50 | 12 | 34.03 | 38.89 | 44.44 | 50.70 | **79.17** | 13.20 | 17.36 | 21.53 | 27.08 | 55.56 |
| 50 | 24 | 31.25 | 34.03 | 36.80 | 44.45 | 73.61 | 10.42 | 11.81 | 14.58 | 22.22 | 50.00 |
| 50 | 36 | 28.47 | 31.94 | 36.11 | 40.28 | 70.83 | 9.03 | 11.11 | 14.58 | 18.06 | 49.30 |
| 50 | 48 | 28.47 | 30.56 | 34.03 | 36.81 | 59.72 | 9.03 | 9.72 | 12.50 | 15.28 | 36.11 |

**Fig. 4.**
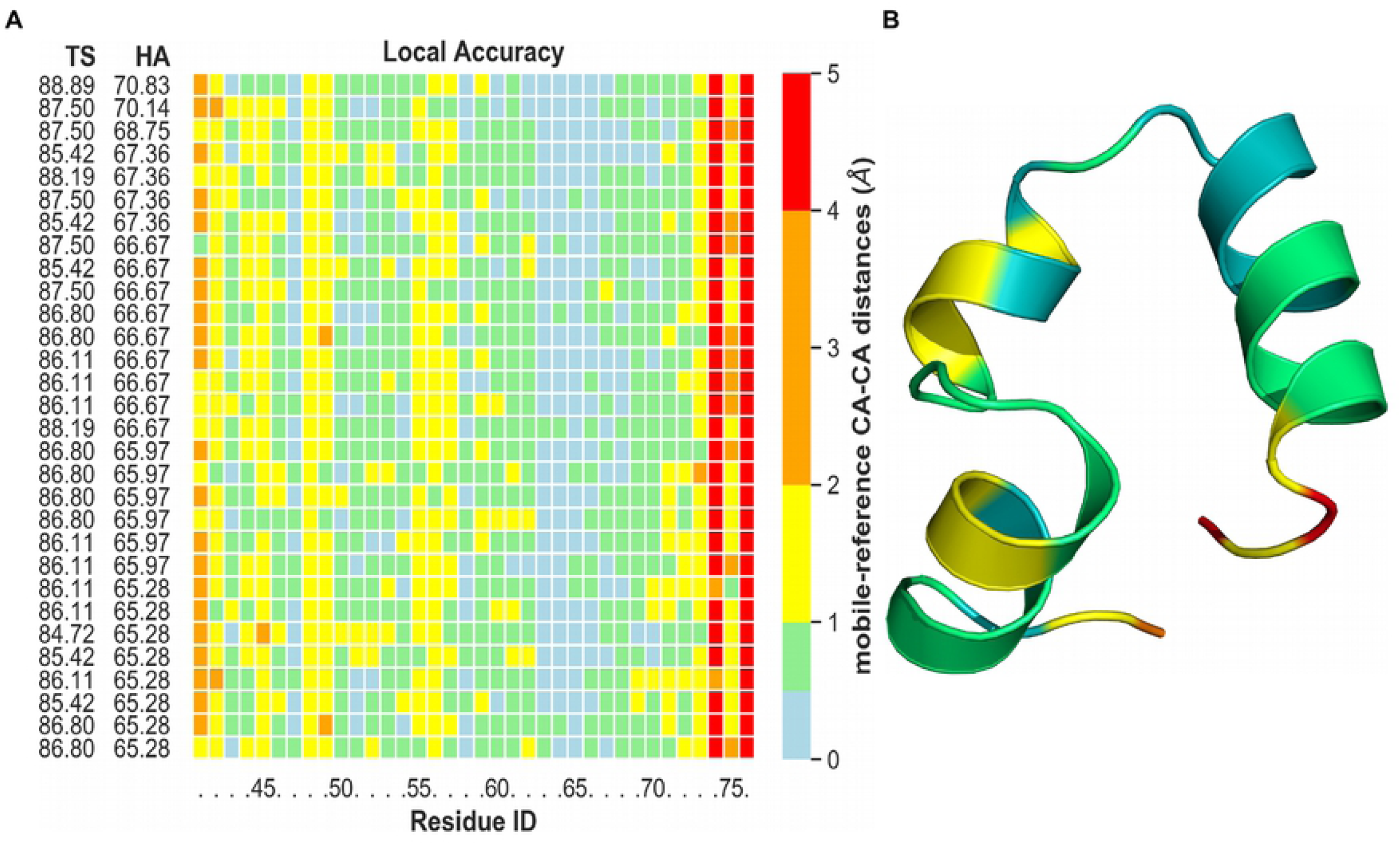
Local accuracy of VHP REX simulation (36 contacts, 75% TPR). (A) Displayed are the best structures ranked by High Accuracy (HA) score and color-coded based on the C_*α*_-C_*α*_ distance to visualize the local accuracy. (B) Best observed tertiary structure of VHP corresponding to the first line of subfigure A.

## Conclusions

Usually, contact information on its own is insufficient to fully determine a protein’s 3D structure. We showed that contact-guided REX MD is capable of delivering physically reasonable structural models of a protein’s conformational state. Contacts derived from various sources can easily be included and thus interpreted in terms of structural ensembles. Our ready-to-use framework for structure determination relies on the well-established and heavily used simulation method of molecular dynamics. As a physics-based method, it is conceptually transparent compared to purely data-driven methods such as AlphaFold [10]. Within one single REX MD run, it is possible to conveniently obtain high-quality results without the need for arduous adjustment of system-specific parameters.

We performed an extensive study on two well-known proteins to test the capabilities and limitations of our framework for protein structure determination. A total of 14 scenarios differing with respect to bias quality by varying true-positive rate and number of used contact restraints were considered. For a facilitated comparison, we kept the coupling strength equal across all restraints. We observed a significant enrichment of native-like conformations as long as the bias quality was above an apparent threshold of 75% TPR. We find that highly error-prone contact information as implemented into 50% TPR scenarios is not sufficient for effective structure determination within REX MD. Typically, it is a priori unknown which of the used contacts are really native, and both experimentally and theoretically derived contact information can mistakenly contain false positives. One possible approach to resolve this issue is a dynamic weighting of contact bias. Initially assigning each contact the same force coefficient, contacts can be monitored regularly and those remaining unrealized can be weakened or completely deactivated accordingly. Furthermore, we evaluated the step-wise improvement by comparing different scenarios with varying numbers of contacts at fixed TPR. In the case of error-free bias, the chance of finding native-like structures increases with the number of contacts included according to our expectations. For more realistic cases with a 75% TPR, we observe significant performance improvements compared to the reference for *L/*2 to *L* restraints, with *L* being the protein sequence length. As a proof of concept, we showed that it is possible to find physically reasonable folds starting from an unfolded state, provided enough turnarounds and a sufficiently long simulation time during REX. To avoid the unnecessary increase of computational costs associated with the large simulation box, one should always start in a pre-estimated folded conformation instead. This would greatly increase the computational performance of each REX run.

Although computationally rather involved, this method is particularly suitable for refinement of often available low-resolution structural models. The underlying force field contains rich information on various physical interactions determining protein dynamics. Since the bias is known and thus can retrospectively be balanced out, the simulated ensemble is thermodynamically correct. Thus, it is in principle possible to infer a free energy landscape with statistical techniques such as the “weighted histogram analysis method”.

## Methods

### Molecular Dynamics

Computer simulations are often used to complement or interpret results of real experiments. Molecular Dynamics (MD) is such an *in silico* approach to study the movements of atoms or biomolecules. Here, one of the many classical force fields is selected and applied to the studied system. The resulting interactions between all atoms are calculated by using Newton’s equations of motions. Typical timesteps are in the order of 1-2 fs. Details of certain mechanisms, such as protein folding or ligand binding, can be observed by analyzing the trajectory of the simulation.

### Replica Exchange

Replica exchange (REX), sometimes referred to as REMD or parallel tempering, is an enhanced sampling technique [20–22]. This efficient method is commonly applied to overcome protein entrapment resulting from the multiple-minima problem during MD simulations. REX simulates *N* non-interacting copies (“replica”) of a system at different temperatures *T*_*i*_. Each replica corresponds to one of *N* MD simulations performed simultanously. After a fixed time *dt* the atom positions and momenta of adjacent replica can be exchanged. The exchange probability is given by the Metropolis criterion [20]

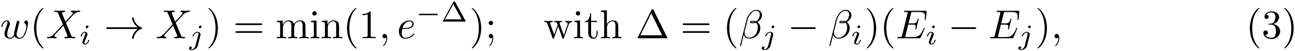

where *X*_*i*_ denotes the state of replica *i*, 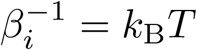 the inverse temperature, and *E*_*i*_ the energy of state *X*_*i*_. Since exchange rates are signifficantly lower for large temperature differences, which can be seen in Δ from Eq (3), it is suffiencent to only exchange adjacent replica.

The intention of REX is to enhance sampling of both high and low energy states. For this reason the temperature range has to be chosen accordingly. Replica at the highest occuring temperatures should have enough energy to overcome potential barriers. Meanwhile, low temperature replica are supposed to explore conformations close to local minima. In combination, REX increases the chance of finding the global energy minimum and thus the native state of a protein. To achieve a random walk it is mandatory to aim for constant exchange rates across all replica and to make sure that all replica are shuffled sufficiently.

### Replica Exchange Temperature Generator

In REX simulations, every replica resembles the dynamics of a canonical ensemble, where the probability distribution of each microstate follows the Boltzmann distribution *e*^*−βE*^. Since exchange rates are propotional to the energy difference of two adjacent replica (see Eq (3)), an exponential temperature distribution is needed to guarantee a random walk in conformation space [13, 20, 28]. This distribution is a priori unknown as it depends on the protein size and number of solvent molecules. However, an initial temperature distribution can be estimated [29]. A simple temperature generator is given by

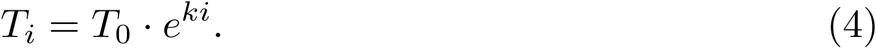

*T*_*i*_ refers to the temperature of replica *i*, while *k* stands for the growth parameter which has to be modified based on the system size. To get more consistent exchange rates during the simulation across all replica, we slightly modified the generator following the equations

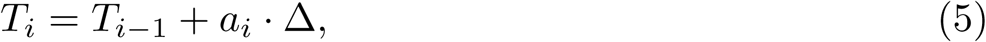

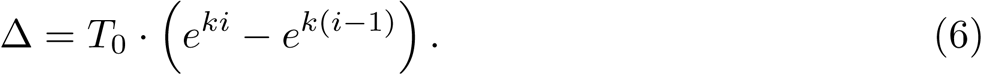

Eq (5) recursively describes the temperature of replica *i*. Δ denotes the temperature difference of two adjacent replica, as specified in Eq (6), while *a*_*i*_ is a step size modifying coefficient. With *a*_*i*_ = 1∀*i* the generator will produce the same temperature distribution as given by Eq (4). To keep the exchange rates almost constant over the whole simulated temperature range, we increased *a*_*i*_ every ten replica by 4%. Used parameters and resulting temperature distributions can be taken from S3 Appendix for Trp-Cage and S4 Appendix for VHP.

### Sigmoid Potential

In order to guide protein folding, we implement an attractive potential to increase the bias towards native-like conformations. In our simplistic model, we apply the force only between the C_*α*_ atoms of used contact pairs. The potential is given by

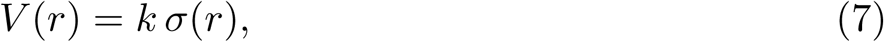

with the force coefficient *k* and the sigmoid function

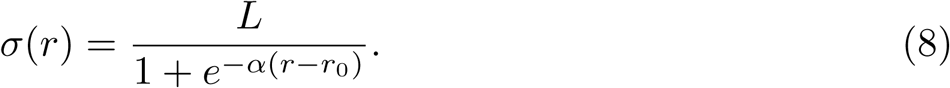

*L* denotes the sigmoid function’s limit, whereas *r* and *r*_0_ stand for the atom distance and equilibirium distance, respectively. Furthermore, *r*_0_ determines the position of the inflection point of the sigmoid function and thus the local maximum of the sigmoid function’s derivative. The parameter *α* affects the S-shape of the function, i.e. how fast the transition from low values to high values takes place. For the sake of simplicity, we set *L* = 1 so that the force only depends on the force coefficient *k*. Fig 5 displays the used sigmoid function *σ*(*r*) in red and its derivative *σ*′(*r*) in green. During all REX MD simulations, we held *k* constant at 10 kJ mol^−1^ (approximately 4 *k*_B_*T* at 300K). Furthermore, we chose *α* = 2.5 nm^*-*1^ and *r*_0_= 1.6 nm.

**Fig. 5.**
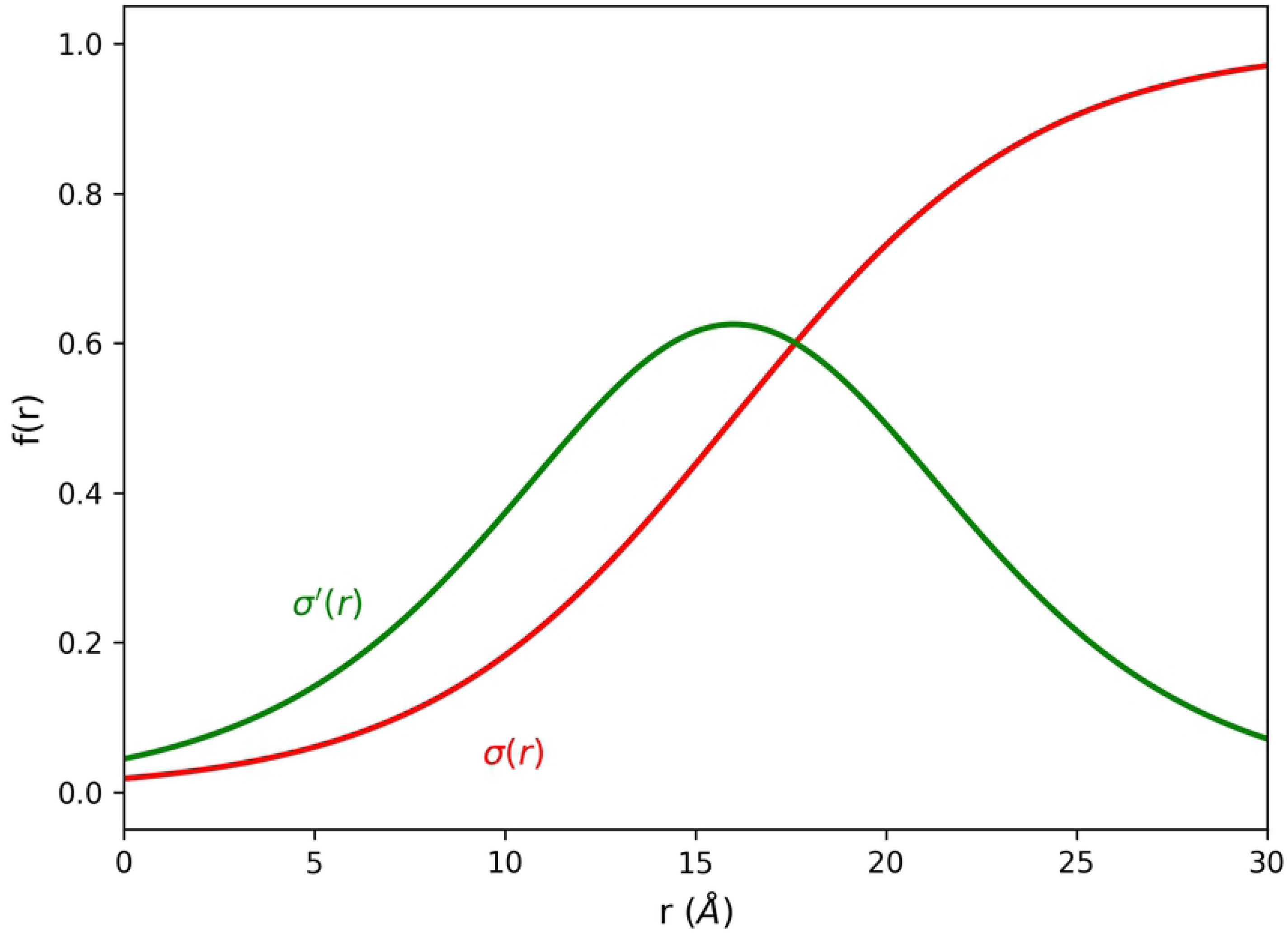
Sigmoid function of bias potential *V* (*r*). The used sigmoid function *σ*(*r*) with parameters *L* = 1, *α* = 2.5 nm^−1^, and *r*_0_= 1.6 nm is represented by the red curve. The derivative *σ*′(*r*) is shown in green.

### Coevolution

The physiological function of a protein is directly linked to its 3D structure determined by the amino acid sequence [30–32]. Through mutation and selection a global structure is preserved within a protein family as a result of coevolution. Destabilizing mutations are statistically compensated by mutations of spatially close amino acids, leaving an evolutionary fingerprint [33, 34]. Coevolution analysis methods such as direct coupling analysis [16] can infer spatially close amino acids based on multiple sequence alignments. The obtained contact pairs can then be used as an additional bias for, e.g., protein folding simulations or structure determination.

### Native Contact Enrichment

One key aspect of this study was to measure the influence and correlation of native and non-native contacts during REX simulations. For this purpose we explicitly used randomly selected contact pairs based on the known structures of the test systems. To quantify the true-positive rate of chosen contact pairs, we define native contacts to fulfill the conditions

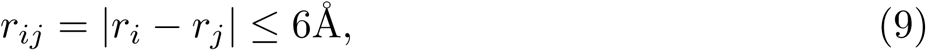

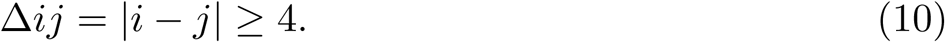

Eq (9) defines a maximum cutoff distance *r*_*ij*_ of 6 Å between C_*α*_ atoms for two residues *i* and *j*. Eq (10) excludes residue pairs which are very close to each other with respect to their sequence position and would appear on the main diagonal of a contact map. Based on these definitions, we created one list with native contacts and another list with non-native contacts. More precisely, we omitted direct neighbors of native contacts within the contact map from the non-native list. For example, if residue pair (*i,j*) is native, then all nine combinations of (*i*′, *j*′) with *i*′ ∈{*i* − 1, *i, i* + 1} and *j*′ ∈{*j* − 1, *j, j* + 1} were excluded. Lastly, we randomly selected contact pairs from each list to construct the different scenarios at fixed TPRs for the method performance study. Contact pairs (*i, j*) used as restraints during the study can be looked up in the contact maps given in S19 Fig to S24 Fig.

### Global Distance Test

During the trajectory analyses backbone RMSD values after structure alignment are considered to initially evaluate the method’s performance. In terms of protein structure determination, however, RMSD is not a proper quantity to assess results as it correlates strongly with the largest displacement between the mobile and target structure. This means if the mobile structure fits the target to a large extent and only one small segment is misaligned, the RMSD becomes disproportionately large. This problem is solved for the so-called Global Distance Test (GDT) [35–37]. Analogously to the RMSD, the mobile structure is first aligned to the target structure. Then the displacement distance of each C_*α*_ residue is calculated and compared with various cutoff thresholds to estimate how similar the two structures are. In a last step, percentages of residues with displacements below a considered threshold are used to calculate score values. The two most common scores are the Total Score (TS),

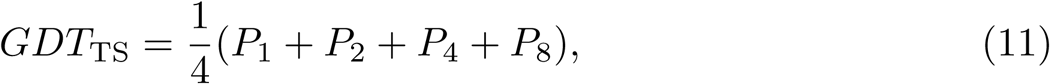

and the High Accuracy (HA) score,

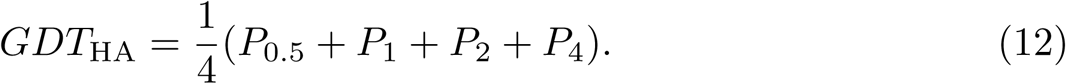

Here, variables *P*_*x*_ denote the percentage of residues with displacements below a distance cutoff of *x* Å. Note that both scores range between 0 and 100 and their interpretation is based on the “fit resolution” set by the applied cutoff distances.

### Setup of REX MD simulations

All simulations were set up on a standard desktop PC using an Intel Core i5-3470 CPU with four cores at 3.20 Ghz frequency. The runs were done with `GROMACS 2016.3` [38, 39] using the `AMBER99SB-ILDN` force field [40] and the `TIP3P` explicit solvent model [41].

Starting from the pdb structure, the protein was unfolded in a normal MD simulation at a high temperature. We selected an unfolded state as initial structure for the REX simulations. All necessary files were generated for the lowest replica temperature *T*_0_. A REX temperature generator based on Eqs (5) and (6) yielded the temperature distribution for *N* replica (cf. S3 Appendix and S4 Appendix). After verifying sufficient exchange rates in short REX simulations (every 1000 steps, rate ≈ 16%), the sigmoid potential based on Eq (7) was provided as a look-up table. Each simulation run for 250*ns* with a stepsize of 2*fs* on 60 (Trp-Cage) and 100 (Villin headpiece) replica. Restraints were added via tabulated bonded interactions [39] to the topology file.

The production runs were performed on the `ForHLR II` cluster. We only used thin nodes, consisting of two Deca-Core Intel Xeon E5-2660 v3 processors (Haswell) with a base clock rate of 2.6 GHz (max. turbo-clock rate of 3.3 GHz), 64 GB main memory, and 480 GB local SSD storage.

## Supporting information

**S1 Appendix. Sample**. **mdp file for REX simulations**.

(PDF)

**S2 Appendix. Look-up table of sigmoid potential**.

(xvg)

**S3 Appendix. Temperature distribution of Trp-Cage REX simulations**.

(PDF)

**S4 Appendix. Temperature distribution of VHP REX simulations**.

(PDF)

**S1 Fig. RMSD Overview during Trp-Cage REX simulation**.

Heatmaps display the backbone RMSD time evolution across all replica. (A) Reference REX simulation without any additional bias. (B) REX simulation with 100% TPR and 6 native contacts as restraints.

(TIF)

**S2 Fig. RMSD Overview during Trp-Cage REX simulation**.

Heatmaps display the backbone RMSD time evolution across all replica. (A) REX simulation with 100% TPR and 12 native contacts as restraints. (B) REX simulation with 100% TPR and 24 native contacts as restraints.

(TIF)

**S3 Fig. RMSD Overview during Trp-Cage REX simulation**.

Heatmaps display the backbone RMSD time evolution across all replica. (A) REX simulation with 100% TPR and 36 native contacts as restraints. (B) REX simulation with 100% TPR and 24 native contacts as restraints.

(TIF)

**S4 Fig. RMSD Overview during Trp-Cage REX simulation**.

Heatmaps display the backbone RMSD time evolution across all replica. (A) REX simulation with 75% TPR and 12 contacts (9 native, 3 non-native) as restraints. (B) REX simulation with 75% TPR and 24 contacts (18 native, 6 non-native) as restraints.

(TIF)

**S5 Fig. RMSD Overview during Trp-Cage REX simulation**.

Heatmaps display the backbone RMSD time evolution across all replica. (A) REX simulation with 75% TPR and 36 contacts (27 native, 9 non-native) as restraints. (B) REX simulation with 75% TPR and 48 contacts (36 native, 12 non-native) as restraints.

(TIF)

**S6 Fig. RMSD Overview during Trp-Cage REX simulation**.

Heatmaps display the backbone RMSD time evolution across all replica. (A) REX simulation with 50% TPR and 12 contacts (6 native, 6 non-native) as restraints. (B) REX simulation with 50% TPR and 24 contacts (12 native, 12 non-native) as restraints.

(TIF)

**S7 Fig. RMSD Overview during Trp-Cage REX simulation**.

Heatmaps display the backbone RMSD time evolution across all replica. (A) REX simulation with 50% TPR and 36 contacts (18 native, 18 non-native) as restraints. (B) REX simulation with 50% TPR and 48 contacts (24 native, 24 non-native) as restraints.

(TIF)

**S8 Fig. RMSD Overview during VHP REX simulation**.

Heatmaps display the backbone RMSD time evolution across all replica. (A) Reference REX simulation without any additional bias. (B) REX simulation with 100% TPR and 6 native contacts as restraints.

(TIF)

**S9 Fig. RMSD Overview during VHP REX simulation**.

Heatmaps display the backbone RMSD time evolution across all replica. (A) REX simulation with 100% TPR and 12 native contacts as restraints. (B) REX simulation with 100% TPR and 24 native contacts as restraints.

(TIF)

**S10 Fig. RMSD Overview during VHP REX simulation**.

Heatmaps display the backbone RMSD time evolution across all replica. (A) REX simulation with 100% TPR and 36 native contacts as restraints. (B) REX simulation with 100% TPR and 24 native contacts as restraints.

(TIF)

**S11 Fig. RMSD Overview during VHP REX simulation**.

Heatmaps display the backbone RMSD time evolution across all replica. (A) REX simulation with 75% TPR and 12 contacts (9 native, 3 non-native) as restraints. (B) REX simulation with 75% TPR and 24 contacts (18 native, 6 non-native) as restraints.

(TIF)

**S12 Fig. RMSD Overview during VHP REX simulation**.

Heatmaps display the backbone RMSD time evolution across all replica. (A) REX simulation with 75% TPR and 36 contacts (27 native, 9 non-native) as restraints. (B) REX simulation with 75% TPR and 48 contacts (36 native, 12 non-native) as restraints.

(TIF)

**S13 Fig. RMSD Overview during VHP REX simulation**.

Heatmaps display the backbone RMSD time evolution across all replica. (A) REX simulation with 50% TPR and 12 contacts (6 native, 6 non-native) as restraints. (B) REX simulation with 50% TPR and 24 contacts (12 native, 12 non-native) as restraints.

(TIF)

**S14 Fig. RMSD Overview during VHP REX simulation**.

Heatmaps display the backbone RMSD time evolution across all replica. (A) REX simulation with 50% TPR and 36 contacts (18 native, 18 non-native) as restraints. (B) REX simulation with 50% TPR and 48 contacts (24 native, 24 non-native) as restraints.

(TIF)

**S15 Fig. Example figure with percentiles**.

Histogram shows the frequencies of occuring Global Distanc Test Total Scores (*GDT*_TS_), during the reference VHP REX simulation. As an example the 80th, 90th and 100th TS percentiles are indicated by red vertical lines.

(TIF)

**S16 Fig. Local accuracy of VHP structures. (reference case)**

(A) Displayed are the best observed structures ranked by High Accuracy (HA) score and color-coded based on the C_*α*_-C_*α*_ distance to visualize the local accuracy. (B) Best observed tertiary structure of VHP corresponding to the first line of subfigure A.

(TIF)

**S17 Fig. Local accuracy of VHP structures. (36 contacts, 100% TPR)**

(A) Displayed are the best observed structures ranked by High Accuracy (HA) score and color-coded based on the C_*α*_-C_*α*_ distance to visualize the local accuracy. (B) Best observed tertiary structure of VHP corresponding to the first line of subfigure A.

(TIF)

**S18 Fig. Local accuracy of VHP structures. (36 contacts, 50% TPR)**

(A) Displayed are the best observed structures ranked by High Accuracy (HA) score and color-coded based on the C_*α*_-C_*α*_ distance to visualize the local accuracy. (B) Best observed tertiary structure of VHP corresponding to the first line of subfigure A.

(TIF)

**S19 Fig. Used restraints during Trp-Cage REX simulations at 100% TPR**.

(A) Contact map displays native contacts as gray squares. Randomly selected contact pairs which were used as restraints are colored based on their batch. (B) Tertiary structure of VHP showing the contact pairs in the same color as in the contact map.

(TIF)

**S20 Fig. Used restraints during Trp-Cage REX simulations at 75% TPR**.

(A) Contact map displays native contacts as gray squares. Randomly selected contact pairs which were used as restraints are colored based on their batch. (B) Tertiary structure of VHP showing the contact pairs in the same color as in the contact map.

(TIF)

**S21 Fig. Used restraints during Trp-Cage REX simulations at 50% TPR**.

(A) Contact map displays native contacts as gray squares. Randomly selected contact pairs which were used as restraints are colored based on their batch. (B) Tertiary structure of VHP showing the contact pairs in the same color as in the contact map.

(TIF)

**S22 Fig. Used restraints during VHP REX simulations at 100% TPR**.

(A) Contact map displays native contacts as gray squares. Randomly selected contact pairs which were used as restraints are colored based on their batch. (B) Tertiary structure of VHP showing the contact pairs in the same color as in the contact map.

(TIF)

**S23 Fig. Used restraints during VHP REX simulations at 75% TPR**.

(A) Contact map displays native contacts as gray squares. Randomly selected contact pairs which were used as restraints are colored based on their batch. (B) Tertiary structure of VHP showing the contact pairs in the same color as in the contact map.

(TIF)

**S24 Fig. Used restraints during VHP REX simulations at 50% TPR**.

(A) Contact map displays native contacts as gray squares. Randomly selected contact pairs which were used as restraints are colored based on their batch. (B) Tertiary structure of VHP showing the contact pairs in the same color as in the contact map.

(TIF)

## Acknowledgments

We thank for computing time on the on the `ForHLR II` cluster, funded by the Ministry of Science, Research and the Arts Baden-Württemberg and by the Federal Ministry of Education and Research. This work is supported by the Helmholtz Association Initiative and Networking Fund under project number ZT-I-0003.

## Author Contributions

**Conceptualization:** Arthur Voronin, Alexander Schug.

**Data Curation:** Arthur Voronin.

**Formal Analysis:** Arthur Voronin.

**Funding Acquisition:** Alexander Schug.

**Investigation:** Arthur Voronin, Alexander Schug.

**Methodology:** Arthur Voronin, Alexander Schug.

**Project Administration:** Alexander Schug.

**Resources:** Arthur Voronin, Alexander Schug.

**Software:** Arthur Voronin.

**Supervision:** Alexander Schug.

**Validation:** Arthur Voronin.

**Visualization:** Arthur Voronin.

**Writing – Original Draft Preparation:** Arthur Voronin, Marie Weiel, Alexander Schug.

**Writing – Review & Editing:** Arthur Voronin, Marie Weiel, Alexander Schug.

## References

1. Guzzo AV. The Influence of Amino Acid Sequence on Protein Structure. Biophysical Journal. 1965;5(6):809–822. doi:10.1016/S0006-3495(65)86753-4.

2. Gan HH, Perlow RA, Roy S, Ko J, Wu M, Huang J, et al. Analysis of protein sequence/structure similarity relationships. Biophysical Journal. 2002;83(5):2781–2791. doi:10.1016/s0006-3495(02)75287-9.

3. Whisstock JC, Lesk AM. Prediction of protein function from protein sequence and structure. Quarterly Reviews of Biophysics. 2003;36(3):307–340. doi:10.1017/S0033583503003901.

4. Benson DA, Cavanaugh M, Clark K, Karsch-Mizrachi I, Lipman DJ, Ostell J, et al. GenBank. Nucleic Acids Research. 2013;41(D1):36–42. doi:10.1093/nar/gks1195.

5. Bateman A. UniProt: A worldwide hub of protein knowledge. Nucleic Acids Research. 2019;47(D1):D506–D515. doi:10.1093/nar/gky1049.

6. Weiel M, Reinartz I, Schug A. Rapid interpretation of small-angle X-ray scattering data. PLoS computational biology. 2019;15(3):e1006900.

7. Reinartz I, Sinner C, Nettels D, Stucki-Buchli B, Stockmar F, Panek PT, et al. Simulation of FRET dyes allows quantitative comparison against experimental data. The Journal of chemical physics. 2018;148(12):123321.

8. Gremer L, Schölzel D, Schenk C, Reinartz E, Labahn J, Ravelli RB, et al. Fibril structure of amyloid-*β*(1–42) by cryo–electron microscopy. Science. 2017;358(6359):116–119.

9. AlQuraishi M. End-to-end differentiable learning of protein structure. Cell systems. 2019;8(4):292–301.

10. Senior AW, Evans R, Jumper J, Kirkpatrick J, Sifre L, Green T, et al. Improved protein structure prediction using potentials from deep learning. Nature. 2020; p. 1–5.

11. Paschek D, Hempel S, García AE. Computing the stability diagram of the Trp-cage miniprotein. Proceedings of the National Academy of Sciences. 2008;105(46):17754–17759.

12. Lindorff-Larsen K, Piana S, Dror RO, Shaw DE. How fast-folding proteins fold. Science. 2011;334(6055):517–520.

13. Hansmann UHE. Parallel tempering algorithm for conformational studies of biological molecules. Chemical Physics Letters. 1997;281(1-3):140–150. doi:10.1016/S0009-2614(97)01198-6.

14. Schug A, Herges T, Wenzel W. Reproducible protein folding with the stochastic tunneling method. Physical review letters. 2003;91(15):158102.

15. Bernardi RC, Melo MC, Schulten K. Enhanced sampling techniques in molecular dynamics simulations of biological systems. Biochimica et Biophysica Acta (BBA)-General Subjects. 2015;1850(5):872–877.

16. Morcos F, Pagnani A, Lunt B, Bertolino A, Marks DS, Sander C, et al. Direct-coupling analysis of residue coevolution captures native contacts across many protein families. Proceedings of the National Academy of Sciences. 2011;108(49):E1293–E1301. doi:10.1073/pnas.1111471108.

17. Schug A, Weigt M, Onuchic JN, Hwa T, Szurmant H. High-resolution protein complexes from integrating genomic information with molecular simulation. Proceedings of the National Academy of Sciences. 2009;106(52):22124–22129. doi:10.1073/pnas.0912100106.

18. Uguzzoni G, John Lovis S, Oteri F, Schug A, Szurmant H, Weigt M. Large-scale identification of coevolution signals across homo-oligomeric protein interfaces by direct coupling analysis. Proceedings of the National Academy of Sciences. 2017;114(13):E2662–E2671. doi:10.1073/pnas.1615068114.

19. De Leonardis E, Lutz B, Ratz S, Cocco S, Monasson R, Schug A, et al. Direct-Coupling Analysis of nucleotide coevolution facilitates RNA secondary and tertiary structure prediction. Nucleic acids research. 2015;43(21):10444–10455.

20. Sugita Y, Okamoto Y. Replica-exchange molecular dynamics method for protein folding. Chemical Physics Letters. 1999;314(1-2):141–151. doi:10.1016/S0009-2614(99)01123-9.

21. Okabe T, Kawata M, Okamoto Y, Mikami M. Replica-exchange Monte Carlo method for the isobaric-isothermal ensemble. Chemical Physics Letters. 2001;335(5-6):435–439. doi:10.1016/S0009-2614(01)00055-0.

22. Sanbonmatsu KY, García AE. Structure of Met-enkephalin in explicit aqueous solution using replica exchange molecular dynamics. Proteins: Structure, Function, and Bioinformatics. 2002;46(2):225–234. doi:10.1002/prot.1167.

23. Neidigh JW, Fesinmeyer RM, Andersen NH. Designing a 20-residue protein. Nature Structural Biology. 2002;9(6):425–430. doi:10.1038/nsb798.

24. Qiu L, Pabit SA, Roitberg AE, Hagen SJ. Smaller and faster: The 20-residue Trp-cage protein folds in 4 *µ*s. Journal of the American Chemical Society. 2002;124(44):12952–12953. doi:10.1021/ja0279141.

25. McKnight CJ, Matsudaira PT, Kim PS. NMR structure of the 35-residue villin headpiece subdomain. Nature Structural Biology. 1997;4(3):180–184. doi:10.1038/nsb0397-180.

26. Freddolino PL, Schulten K. Common structural transitions in explicit-solvent simulations of villin headpiece folding. Biophysical Journal. 2009;97(8):2338–2347. doi:10.1016/j.bpj.2009.08.012.

27. Lee H, Turilli M, Jha S, Bhowmik D, Ma H, Ramanathan A. DeepDriveMD: Deep-Learning Driven Adaptive Molecular Simulations for Protein Folding. 2019 IEEE/ACM Third Workshop on Deep Learning on Supercomputers (DLS). 2019; p. 12–19. doi:10.1109/dls49591.2019.00007.

28. Sindhikara D, Meng Y, Roitberg AE. Exchange frequency in replica exchange molecular dynamics. Journal of Chemical Physics. 2008;128(2):024103. doi:10.1063/1.2816560.

29. Patriksson A, van der Spoel D. A temperature predictor for parallel tempering simulations. Physical Chemistry Chemical Physics. 2008;10(15):2073. doi:10.1039/b716554d.

30. Anfinsen CB. Principles that Govern the Folding of Protein Chains. Science. 1973;181(4096):223–230. doi:10.1126/science.181.4096.223.

31. Sadowski MI, Jones DT. The sequence-structure relationship and protein function prediction. Current Opinion in Structural Biology. 2009;19(3):357–362. doi:10.1016/j.sbi.2009.03.008.

32. Bepler T, Berger B. Learning protein sequence embeddings using information from structure. 7th International Conference on Learning Representations, ICLR 2019. 2019; p. 1–17.

33. Sutto L, Marsili S, Valencia A, Gervasio FL. From residue coevolution to protein conformational ensembles and functional dynamics. Proceedings of the National Academy of Sciences of the United States of America. 2015;112(44):13567–13572. doi:10.1073/pnas.1508584112.

34. Szurmant H, Weigt M. Inter-residue, inter-protein and inter-family coevolution: bridging the scales. Current Opinion in Structural Biology. 2018;50:26–32. doi:10.1016/j.sbi.2017.10.014.

35. Zemla A. LGA: A method for finding 3D similarities in protein structures. Nucleic Acids Research. 2003;31(13):3370–3374. doi:10.1093/nar/gkg571.

36. Kufareva I, Abagyan R. Methods of protein structure comparison. In: Homology Modeling. Springer; 2011. p. 231–257.

37. Modi V, Xu Q, Adhikari S, Dunbrack RL. Assessment of template-based modeling of protein structure in CASP11. Proteins: Structure, Function, and Bioinformatics. 2016;84(1):200–220. doi:10.1002/prot.25049.

38. Van Der Spoel D, Lindahl E, Hess B, Groenhof G, Mark AE, Berendsen HJC. GROMACS: Fast, flexible, and free. Journal of Computational Chemistry. 2005;26(16):1701–1718. doi:10.1002/jcc.20291.

39. Abraham MJ, van der Spoel D, Lindahl E, Hess B, the GROMACS development team. GROMACS User Manual version 2016.3; 2017. Available from: www.gromacs.org.

40. Lindorff-Larsen K, Piana S, Palmo K, Maragakis P, Klepeis JL, Dror RO, et al. Improved side-chain torsion potentials for the Amber ff99SB protein force field. Proteins: Structure, Function and Bioinformatics. 2010;78(8):1950–1958. doi:10.1002/prot.22711.

41. Jorgensen WL, Chandrasekhar J, Madura JD, Impey RW, Klein ML. Comparison of simple potential functions for simulating liquid water. The Journal of Chemical Physics. 1983;79(2):926–935. doi:10.1063/1.445869.

